# Intracellular pools of DAG-activated TRPC3 channels are essential for TLR4 activation

**DOI:** 10.1101/2021.02.24.432657

**Authors:** Javier Casas, Clara Meana, José Ramón López-López, Jesús Balsinde, María A. Balboa

## Abstract

Toll-like receptor 4, the receptor for bacterial lipopolysaccharide (LPS), drives inflammatory responses that protect against pathogens and boost the adaptive immunity. LPS responses are known to depend on calcium fluxes, but the molecular mechanisms involved are poorly understood. Here we present evidence that the transient receptor potential canonical channel 3 (TRPC3) is activated intracellularly during macrophage exposure to LPS and is essential for Ca^2+^ release from internal stores. In this way TRPC3 participates in cytosolic Ca^2+^ elevations, TLR4 endocytosis, activation of inflammatory transcription factors and cytokine upregulation. We also report that TRPC3 is activated by diacylglycerol (DAG) generated by the phosphatidic acid phosphatase lipin-1. In accord with this, lipin-1-deficient cells show reduced Ca^2+^ responses to LPS challenge. A cameleon indicator directed to the endoplasmic reticulum shows that this is the organelle from which TRPC3 mediates the Ca^2+^ release. Finally, pharmacological inhibition of TRPC3 reduces systemic inflammation induced by LPS in mice. Collectively, our study unveils a central component of LPS-triggered Ca^2+^ signaling that involves intracellular sensing of lipin-1-derived DAG by TRPC3.

## INTRODUCTION

Receptors on the surface of innate immune cells recognize multiple molecules derived from pathogens but also molecules generated during a cellular insult that act as danger signals. These receptors initiate an inflammatory reaction that helps to eliminate the pathogen and repair the damaged area. TLR4, a prominent member of the Toll-like receptor family, is the canonical receptor for lipopolysaccharide (LPS) from gramnegative bacteria ^1^. LPS recognition by TLR4 initiates a cascade of events that culminates in the activation of the transcription factors AP-1, NF-κB, or IRF-3 through different signaling branches ^2–5^. The final outcome is the transcriptional upregulation of cytokines, enzymes and proteins that help inactivate the invading microorganism. However, excessive TLR4 activation orchestrated by elevated concentrations of LPS could be detrimental and constitutes the basis of some serious diseases such as sepsis. Hence, a finer knowledge of TLR4-mediated signaling is key to find helpful targets for the management of infections and inflammatory diseases.

It is now clearly established that complete TLR4 responses require cellular Ca^2+^ fluxes and [Ca^2+^]i elevations ^6–10^. Ca^2+^ signaling in immune cells has traditionally been considered to be due to the release of Ca^2+^ from internal stores, such as the endoplasmic reticulum (ER), followed by Ca^2+^ influx through plasma membrane channels to restore homeostasis ^11^. This mechanism is known as store-operated Ca^2+^ entry (SOCE). Regarding TLR4 associated Ca^2+^ fluxes, it has been proposed that they are initiated by the activation of phospholipase Cγ_2_ (PLCγ_2_) at the plasma membrane, mediated by the TLR4 co-receptor CD14 ^7,9^. While not directly demonstrated, this model assumes that PLCγ_2_–derived inositol 1,4,5-trisphosphate (IP_3_) interacts with receptors at the ER which mediate Ca^2+^ release. As a consequence, SOCE mechanisms are activated and participate in the [Ca^2+^]i rise. The discovery that elimination of the proteins that regulate SOCE trough interaction with ORAI channels at the plasma membrane, i.e. STIM1 and STIM2, has no discernible effect on LPS signaling in macrophages ^12^, has encouraged very active research programs to identify alternative players in Ca^2+^ dynamics. An important aspect to consider is that elimination of TLR4 or its co-receptor CD14, decreases the capacity of LPS to induce a rise in [Ca^2+^]i ^10^. Hence, Ca^2+^ fluxes and the channels that mediate them, have to be directly regulated by LPS-generated signals.

Some members of the transient receptor potential canonical (TRPC) channel superfamily can be activated by second messengers of lipid nature. TRPC3, together with TRPC6 and TRPC7, are calcium-permeable nonselective cation channels that share the special attribute of being activated by diacylglycerol (DAG) ^13,14^. Of these, TRPC3 is the best characterized in innate immune cells ^15^. TRPC3, was initially described to be involved in SOCE^16^. However, this channel has the capacity to directly bind DAG thanks to a lateral fenestration in its pore domain which is critical for channel gating ^17,18^. It is thought that DAG binding to TRPC3 constitutes the key event that drives TRPC3 activation and channel-mediated Ca^2+^ signaling. DAG may be transiently generated during signaling through the action of PLCs on membrane phosphoinositides, but it may also be produced in a long lasting manner by other means, involving the concerted action of several enzymes and pathways ^19^. In this regard, we have recently described the generation of DAG during TLR4 by lipin-1 ^20^. Lipin-1 is a prominent member of a family of phosphatidic acid (PA) phosphatase enzymes with key roles in metabolism and signaling ^21–23^. In the absence of lipin-1 many TLR4 signaling events are altered in macrophages, including cytokine production and inflammation in animal models of disease ^20,24^. Thus, lipin-1-derived DAG is of great importance during LPS challenge albeit its downstream direct targets are poorly defined.

In this work we have investigated the possible role of TRPC3 during TLR4 stimulation in human macrophages. By using live cell microscopy, DAG probes specifically targeted to different cellular membranes, and cameleon ER-targeted indicators, we have found that TRPC3 participates in the Ca^2+^ fluxes that follow LPS activation of macrophages. Unanticipatedly, we find that the channel is activated inside the cell, close to where lipin-1 generates the DAG. Further, we establish the importance of these events for TLR4 internalization, signal transduction and cytokine production. We also describe that TRPC3 specific inhibitors ameliorate inflammation in mice. Due to the recent development of photopharmacology and optochemical genetics of TRPC3 that could be the foundation for novel therapeutics ^25,26^ these discoveries highlight pharmacological targeting of TRPC3 as an attractive new strategy for the treatment of sepsis and inflammatory diseases.

## RESULTS

### Involvement of TRPC3 channels in TLR4 signaling leading to cytokine production

We began this study by analyzing the expression of DAG-regulated TRPC channels in human THP-1 macrophages by using conventional semi-quantitative PCR. As shown in Figure 1, TRPC3, 6 and 7 were expressed in these cells, with TRPC3 exhibiting the highest expression levels (**Fig. 1a**). To analyze whether this channel participates in the inflammatory activation of human macrophages via TLR4 receptors, we utilized cells with deficient expression of TRPC3 by siRNA silencing. Reduced expression levels of TRCP3 (**Fig. 1b**) strongly blunted the LPS-induced upregulation of a number of proinflammatory genes, including *COX2, TNFA, IL1B* and *IL6* (**Fig. 1c**). Overexpression of the TRPC3 channel in engineered HEK cells to express the human sequences for TLR4/MD2/CD14 (HEK-TLR4 cells) (**Figs. 1d and 1e**), markedly increased the expression levels of *COX2, TNFA, IL1B, IL12B* and *IL23A* after cell treatment with LPS (**Fig. 1e**). Control HEK-TLR4 cells not overexpressing TRPC3 were unable to induce the expression of proinflammatory genes, an effect which is likely due to the absence of serum in the incubation media ^27^. In contrast to TRPC3, overexpression of TRPC6 did not have any effect on the expression of proinflammatory genes upon LPS stimulation (**Figs 1d and 1e**). Collectively, these results support a key effector role for TRPC3 in LPS-induced proinflammatory gene expression.

**Figure 1.**
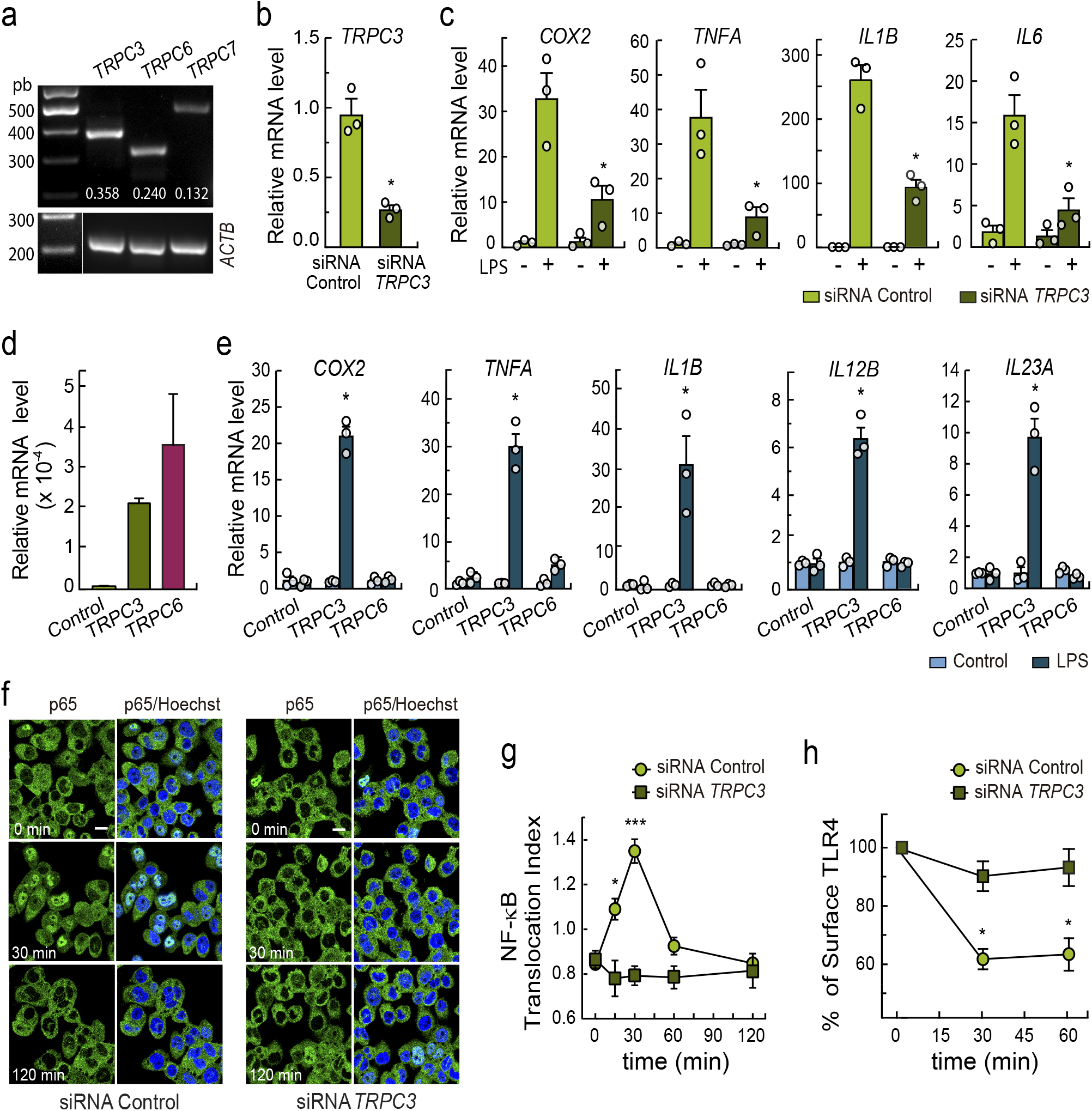
TRPC3 is required for LPS mediated upregulation of proinflammatory genes and signaling. **(a)** Semi-quantitative analysis of the expression of DAGsensitive TRPC channels in THP-1 macrophages. The image is representative of 3 independent experiments. Data quantification using *ACTB* for normalization is shown in white. **(b)** THP-1 macrophages were silenced with control siRNAs or specific siRNAs for *TRPC3*. RT-qPCR quantification of *TRPC3* mRNA levels is shown. Error bars represent SEM (n=3). **(c)** Silenced THP-1 macrophages were treated with 100 ng/ml LPS for 5 h and mRNA levels were quantified by RT-qPCR. Changes in mRNA levels relative to untreated cells are represented. Error bars represent SEM (n=3). **(d)** HEK-TLR4 cells were transfected with TRPC3, TRPC6 o empty plasmids (control cells). Changes in mRNA levels relative to control cells are shown. **(e)** Cells treated as in d were stimulated with 1 μg/ml LPS for 3 h, and mRNA levels analyzed by RT-qPCR. Changes in mRNA levels relative to control cells are shown. Error bars represent SEM (n=3). **(f, g)** THP-1 macrophages were silenced with control siRNAs or specific siRNAs for *TRPC3*, treated with 100 ng/ml LPS for the indicated periods of time and immunostained using antibodies against p65 NF-κB. Nuclei were counterstained with Hoechst 33342. Confocal microscopy pictures **(f)** and nuclear/cytosolic ratios of p65 fluorescence **(g)** are shown. Error bars represent SEM (n=3 images with 75-100 cells/image). Scale bar represents 10 μm. **(h)** Flow cytometry analysis of cell surface TLR4 performed in silenced THP-1 macrophages treated with 100 ng/ml LPS for the indicated periods of time. Data are shown are the relative percentage of cell surface TLR4 calculated as described in Materials and Methods. *, p < 0.05; ***, p < 0.001, by Student’s t test.

NF-κB is a transcription factor implicated in the upregulation of many of the genes described above, and its full activation depends on Ca^2+^ fluxes ^8,10,28,29^. Thus, we studied next the activation of this transcription factor by analyzing the translocation to the nucleus of one of its classical subunits, p65. LPS treatment of the THP-1 macrophages induced a time-dependent translocation of p65 to the nuclear compartment. Such translocation reached a maximum at 30 min after LPS exposure (**Figs. 1f, and 1g**). In contrast, TRPC3-deficient macrophages showed no translocation of p65.

Activation of the MAPK cascade is another hallmark of cell activation via TLR4 ^30^. Some of the MAPKs are known to participate in the activation of the transcription factor AP-1, which assists in the upregulation of inflammatory genes ^30^. To evaluate whether TRPC3 was involved in LPS-induced MAPK activation, we blocked the activity of the channel by using the specific cell-permeable inhibitor Pyr10 ^31^. THP-1 macrophages treated with the inhibitor exhibited lower phosphorylation levels for p38 and ERK (p42/p44) after LPS stimulation. However, JNK (p46/p54) was not affected (**Fig. S1**).

In view of the above results we hypothesized that TRPC3 could be implicated in an early event following engagement of TLR4. One of these is the internalization of TLR4 itself by endocytosis ^32,33^. Endosomal TLR4 forms complexes with TRAM-TRIF, which initiates a cascade of events culminating in the activation of the transcription factor IRF3, and amplification of NF-κB activation ^34^. Thus we evaluated TLR4 levels at the cell membrane of macrophages by flow cytometry. **Fig. 1h** shows that TRPC3-deficient macrophages exhibited much greater labeling of membrane TLR4 than control cells during LPS activation. These data reveal defective internalization in cells lacking TRPC3, and are fully consistent with the cells exhibiting decreased activation of NF-κB, as previously shown. Altogether, the results suggest a key involvement for the TRPC3 channel in TLR4-mediated cell activation and consequently, in the upregulation of inflammatory genes by human macrophages.

### LPS-induced Calcium Signaling via TLR4 is Mediated by TRPC3

Elevations in intracellular calcium ([Ca^2+^]i) during TLR4 activation constitute a required event for both endocytosis and signaling ^7,32,33^. In the next series of experiments we used live-cell imaging to explore the involvement of TRPC3 in the regulation of Ca^2+^ fluxes during TLR4 engagement. Using the dynamic single-wavelength fluorescent Ca^2+^ indicator Fluo-4, whose increase in fluorescence intensity reflects a rise in the cytoplasmic Ca^2+^ level, we detected significant [Ca^2+^]i increases after cellular treatment with LPS (**Fig. 2a, 2b**), in agreement with previous reports ^6,9,10^. Interestingly, macrophages deficient in TRPC3 exhibited smaller [Ca^2+^]i rises than control cells (**Fig. 2b**). Quantification of the areas under the curves demonstrated that, in the absence of TRPC3, cells experienced a reduction in [Ca^2+^]i of ~60% during activation. To further assess the role of TRPC3, we used the cell-permeable selective blocker Pyr10, and observed that acute inhibition of TRPC3 profoundly reduced the [Ca^2+^]i rise in macrophages (**Fig. 2c**). From these data we anticipated that, as a consequence of the effect on Ca^2+^ fluxes, Pyr10 would reduce inflammatory gene expression. **Figure 2d** shows that this was indeed the case; the expression of *COX2, TNFA, IL6, IL12B* and *IL23* was markedly reduced in the presence of Pyr10.

**Figure 2.**
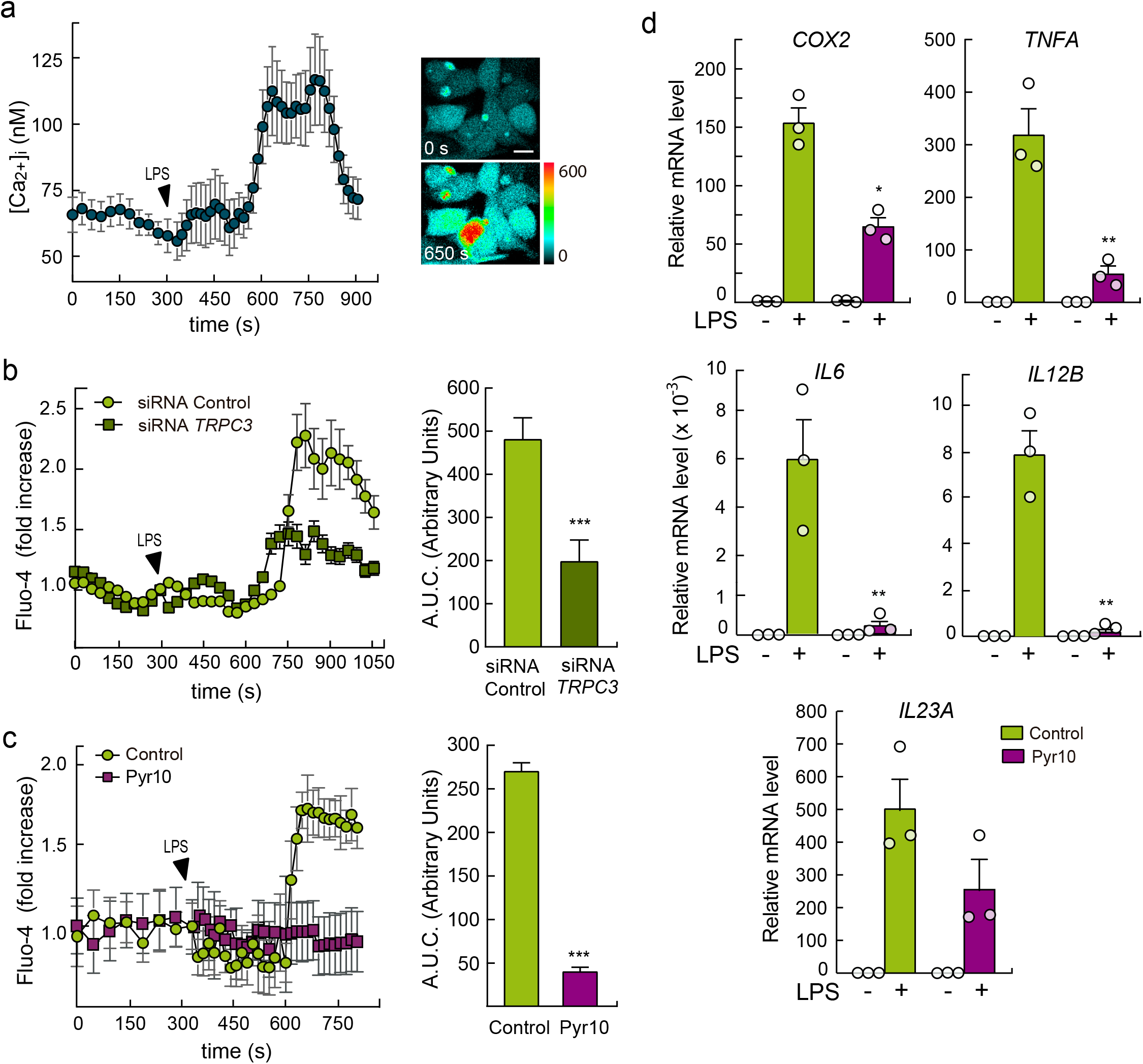
LPS-induced Ca^2+^ fluxes depend on TRPC3. **(a)** THP-1 macrophages were labeled with Fluo-4, and fluorescence was recorded before and after treating the cells with 100 ng/ml LPS, as indicated. Absolute changes in [Ca^2+^]i are shown (n = 25 cells) (left panel). Representative pictures showing fluorescence intensities at 0 s and 350 s after LPS treatment are shown (right panel). Scale bar represents 10 μm. **(b)** THP-1 macrophages silenced with siRNA control or against *TRPC3* were treated as in a. Relative changes in [Ca^2+^]i are shown (left panel). Quantification of the area under the curve (AUC) is shown in arbitrary units (right panel). Error bars represent SEM (siRNA Control, n=28; siRNA *TRPC3*, n=23). ***, p < 0.001, Student’s t test. **(c)** Macrophages were treated as in a and changes in [Ca^2+^]i were analyzed in the absence or presence of 10 μM Pyr10 (left panel). Quantification of the area under the curve (AUC) is shown in arbitrary units (right panel). Error bars represent SEM (n= 25) ***, p < 0.001, Student’s t test. **(d)** THP-1 macrophages were treated with 100 ng/ml LPS for 5 h in the absence or presence of 10 μM Pyr10. mRNA levels for the indicated genes where analyzed by RT-qPCR. Error bars represent SEM (n=3). *, p < 0.05; **, p < 0.01; Student’s t test.

Collectively, these data indicate that TRPC3 participates in Ca^2+^ fluxes that occur during TLR4 activation in macrophages and that this event is important for the production of inflammatory cytokines.

### Non-Plasma Membrane DAG Drives TRPC3-dependent Inflammatory Activation

To investigate further the relationship between TRPC3 and TLR4 activation, we carried out whole-cell patch-clamp recordings. The advantage of this technology is that it can unequivocally, measure the activation of channels located in the plasma membrane. Based on the studies measuring [Ca^2+^]i (**Fig. 2**) and in the biology of the TRPC3 channel ^26^ we anticipated that a significant inward current during treatment of the cells with LPS would be detected, reflecting the activation of ion currents through the channel. However, we failed to detect such activation (**Figs. 3a and 3b**). Because TRPC3 has been described as a DAG-activated channel ^13^, we treated macrophages with the synthetic DAG 1-oleyl-2-acetyl-*sn*-glycerol (OAG). As expected, upon addition of OAG we did detect the activation of a Pyr10 sensitive current (**Figs. 3a and 3b**). These results clearly show that macrophages express channels (TRPCs) whose currents can be activated by DAG but, interestingly, those channels are not sensitive to LPS, at least at the plasma membrane.

**Figure 3.**
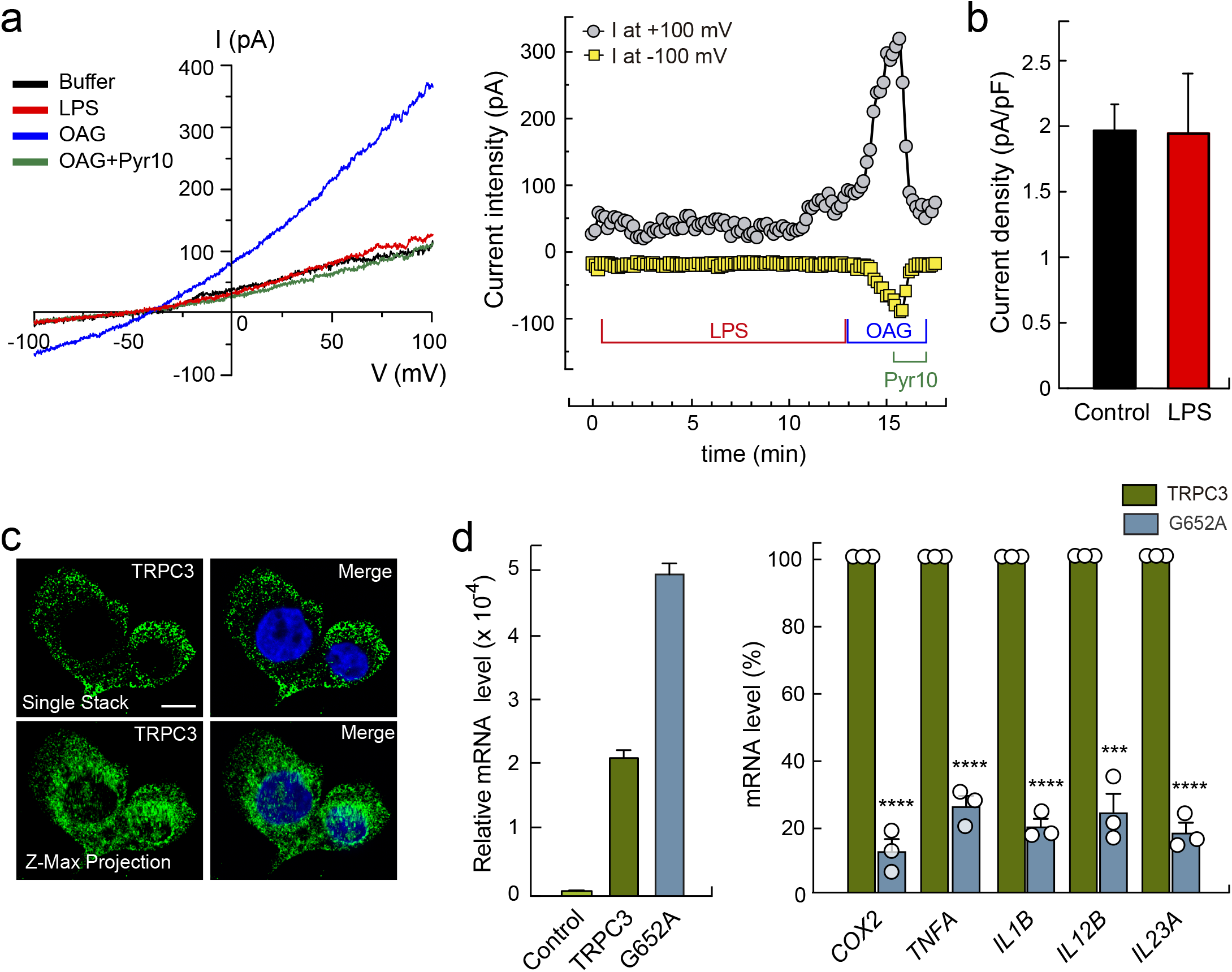
DAG drives TRPC3-dependent inflammatory activation not at the plasma membrane. **(a)** Typical recordings of whole-cell currents obtained at −100 and +100 mV in a peritoneal macrophage. Left panel shows the full current to voltage relationship obtained with the ramps applied to cells treated with 200 ng/ml LPS, 10 μM OAG, or 10 μM OAG plus 10 μM Pyr10 as indicated. Right panel shows time courses of current densities at the indicated conditions. **(b)** Average Pyr10 sensitive current densities obtained at −100 mV after 10 minutes of stimulation with 200 ng/ml LPS (n=5 cells) or after 10 minutes of recording in control solution (n=5 cells). **(c)** THP-1 macrophages were immunostained with antibodies against TRPC3 (green) and counterstained with Hoechst 333342 (nuclei, blue). Images show analyses of fluorescence by confocal microscopy (top). Images of maximum z-projection analyses of 22 stacks are also shown (bottom). Scale bar represents 10 μm. **(d)** HEK-TLR4 cells were transfected with wild type TRPC3, the mutant G652A, o empty plasmids (control cells). mRNA levels of *TRPC3* relative to control cells are represented (left panel). Cells were then treated with 1 μg/ml LPS for 3 h. mRNA levels for the indicated genes (right panel) where analyzed by RT-qPCR and mRNA changes in cells overexpressing the mutant G652A are expressed as % relative to wild type TRPC3-expressing cells. Error bars represent SEM (n=3). ***, p < 0.001; ****, p < 0.0001, Student’s t test.

Although TRPC3 channels are localized in the plasma membrane ^35^, they may also be present in intracellular membranes, particularly those of the endoplasmic reticulum ^36^ and mitochondria ^37^. Using specific TRPC3 antibodies, we performed immunocytochemistry experiments in human macrophages to establish the localization of TRPC channel in these cells. As shown in **Fig. 3**, TRPC3 was mostly found in cytoplasm membranes, being especially concentrated in membranes close to the nucleus. This was particularly evident when all the images obtained from single stacks were subjected to z-maximum projection (**Fig. 3c**).

The results of the patch-clamp experiments, together with the finding that macrophages exhibit much of their TRPC3 content at intracellular locations, moved us to study whether DAG was required or not for TRPC3 to regulate inflammatory actions mediated by TLR4. To this end, we took advantage of the recent finding that a single mutation in TRPC3, G652A, modifies its capacity to recognize DAG, resulting in loss of function ^17^. We transfected cells with the G652A TRPC3 mutant and studied the capacity of the cells to produce inflammatory effectors. The results indicated a dramatic difference between cells transfected with wild type TRPC3 and the G652A mutant regarding their capacity to upregulate *COX2, TNFA, IL1B, IL12B and IL23A* after LPS stimulation (**Fig. 3d**). Note that, in these experiments, the cells expressed the mutant G652A TRPC3 at higher levels than wild type TRPC3. Collectively, these results suggest that DAG recognition by TRPC3 is key for activation of inflammatory responses to TLR4 receptor stimulation.

### LPS Increases DAG Levels in the ER

To assess whether TRPC3 could be activated by DAG in endomembranes, we analyzed physiological DAG dynamics during TLR4 stimulation. We used genetically encoded FRET-based DAG biosensors that can be targeted specifically to different biological membranes: namely Daglas-pm1 and Daglas-em1. Daglas-pm is directed to the plasma membrane through an 11 amino-acid sequence of N-Ras that allows its prenylation ^38,39^ Daglas-em1, directed to ER membranes, is obtained by mutating the C181 of N-Ras to Ser ^39^. Transfected biosensors localized as expected in both THP-1 macrophages and HEK-TLR4 cells (**Figs. 4a and 4b**). Cells expressing Daglas-pm1 immediately showed a substantial increase in intramolecular FRET/CFP ratio when stimulated with the DAG analog, phorbol 12-myristate 13-acetate (PMA) (**Fig. 4a**). However, when the cells were stimulated with LPS, no substantial effect on the fluorescence emission ratio was detected, suggesting that this particular sensor is not detecting changes in DAG levels at the plasma membrane under these conditions. In contrast, cells expressing the ER-directed Daglas-em1 sensor, showed a significant increase in FRET/CFP ratio after LPS stimulation in both THP-1 macrophages and HEK-TLR4 cells (**Fig. 4b**).

**Figure 4.**
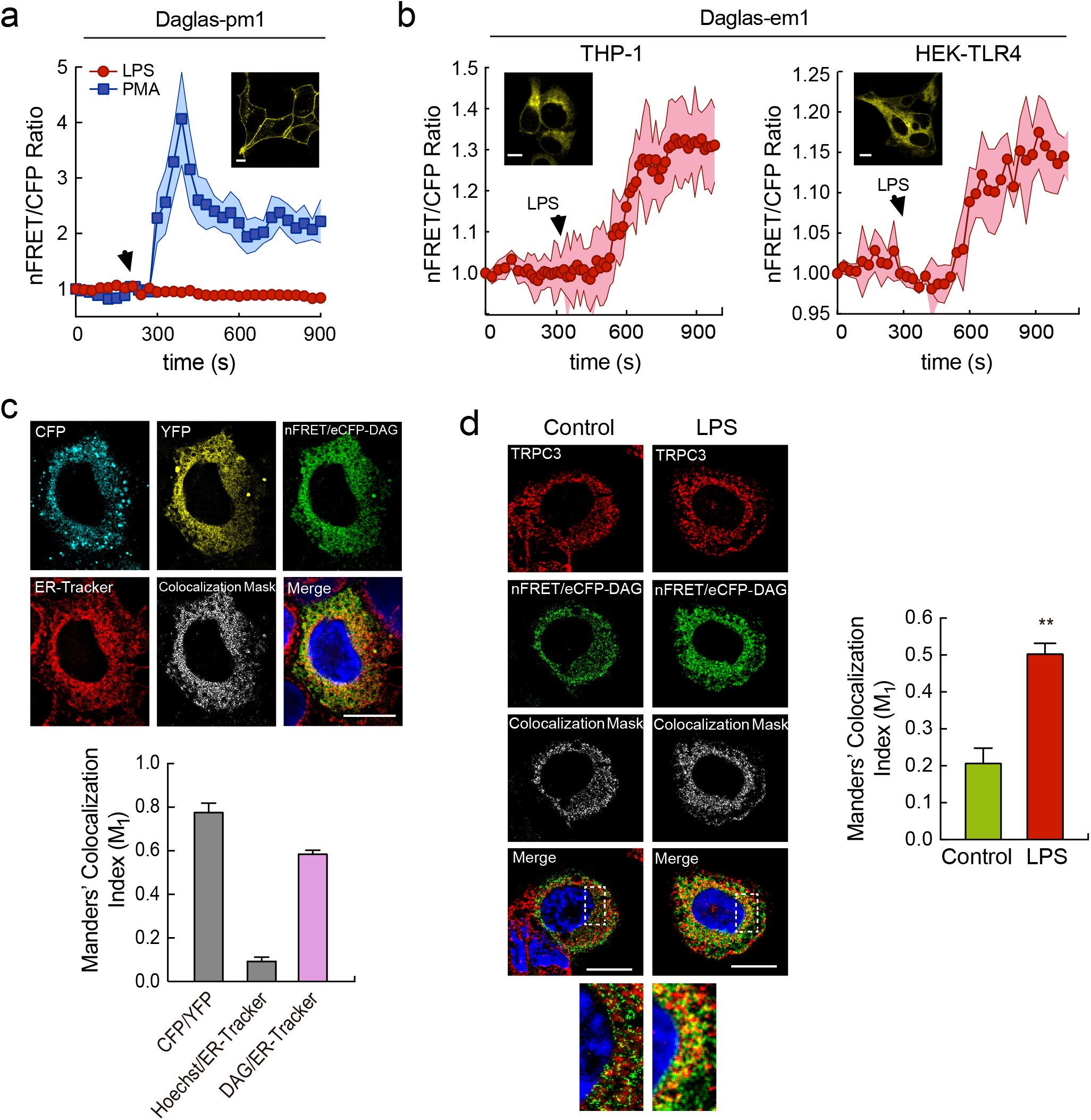
TRPC3 colocalizes with DAG at the ER. **(a)** HEK-TLR4 cells transfected with the sensor Daglas-pm1 were stimulated with 1 μg/ml LPS (red circles) or 1 μM PMA (blue squares), as indicated. Time course changes in nFRET/CFP ratio are shown. Insert show representative images of sensor fluorescence. Scale bar represents 10 μm. **(b)** THP-1 macrophages (left panel) and HEK-TLR4 cells (right panel), transfected with the sensor Daglas-em1, were stimulated with 100 ng/ml and 1 μg/ml LPS respectively. Time course of the changes in nFRET/CFP ratio are shown. Shadows in pink represent SEM (n=27). Inserts show representative images of sensor fluorescence. Scale bar represents 10 μm. **(c)** THP-1 macrophages transfected with Daglas-em1 sensor were stimulated with 100 ng/ml LPS for 10 min and stained with 200 nM ER-Tracker and Hoechst. Fluorescence from CFP (cyan), YPF (yellow), nFRET/CFP (DAG, green), Hoechst (nuclei, blue), ER-Tracker (ER, red), was analyzed by confocal microscopy and images, including merge and the colocalization mask between DAG and ER-Tracker fluorescences (white), are shown (upper panel). Scale bar represents 10 μm. Manders’ colocalization indexes (m1) are shown (bottom panel). Error bars represent SEM (n=28). **(d)** THP-1 macrophages, transfected with the Daglas-em1 sensor, were treated or not with 100 ng/ml LPS for 5 min, fixed and stained with antibodies against TRPC3 (red) and Hoechst (blue). Fluorescence images analyzed by confocal microscopy are shown. Images on the bottom are a detailed amplification of the framed zones from the merge images (left panel). Scale bar represents 10 μm. Manders’ colocalization indexes (m1) between DAG (nFRET/CFP, green) and TRPC3 (red) in cells stimulated or not with LPS are shown (right panel). Error bars represent SEM (n=22 cells). **, p < 0.01; Student’s t test.

To precisely identify the endomembranes where the relevant DAG pool accumulates during TLR4 stimulation, THP-1 macrophages were transfected with the Daglas-em1 biosensor, stimulated with LPS, and subsequently labeled with the ER specific marker ER-Tracker-TR (**Fig. 4c**). Colocalization between DAG and ER was studied by using the Manders’ colocalization index m1 ^40,41^. The colocalization index between FRET/CFP (representing DAG content) and ER-Tracker-TR (representing ER) was 0.58 ± 0.02. Given that the colocalization index between CFP and YFP –which were engineered into the same molecule–, was 0.78 ± 0.04, these data suggest that a substantial amount of the DAG generated during macrophage stimulation with LPS accumulates in the ER.

### TRPC3 Colocalizes with DAG at the ER and Regulates [Ca^2+^]er during TLR4 activation

We investigated next whether the DAG generated in endomembranes upon LPS treatment localized within the same intracellular structures as TRPC3. Macrophages expressing the sensor Daglas-em1 were stimulated with LPS, immunostained with antibodies against TRPC3, and the colocalization between DAG and TRPC3 was analyzed. A marked increase in the colocalization index in stimulated cells compared with control cells in areas adjacent to the nucleus, where the ER is located, was appreciated (**Fig. 4d**). These results suggest that a significant amount of TRPC3 is found at the same structures where DAG accumulates, i.e. ER membranes, during LPS treatment.

Based on the findings that: (i) ion currents through TRPC3 are not activated at the plasma membrane by LPS, (ii) the channel mediates Ca^2+^ fluxes during TLR4 activation, and (iii) the channel is present mainly in intracellular membranes, we reasoned that TRPC3 could be involved in regulating the movement of Ca^2+^ from intracellular stores. To investigate this possibility, we took advantage of a FRET-based calcium “cameleon” indicator engineered to reside in the lumen of the ER (vCY4er) ^42^. This indicator detects changes in [Ca^2+^]er during cellular stimulation ^42^. Cells transfected with this indicator showed a reticular pattern of fluorescence characteristic of ER localization, as initially described ^42^, and underwent a decrease in FRET/eCFP ratio when Ca^2+^ decreased in the organelle due to thapsigargin treatment (**Fig. 5a**). LPS treatment reduced the FRET/eCFP ratio of the cameleon indicator (**Fig. 5b**). Interestingly, the effect was almost abolished when TRPC3-deficient macrophages were used (**Fig. 5b**), and also in the presence of Pyr10 (**Fig. 5c**). These fluorescence changes reflect a reduction in [Ca^2+^]er during stimulation, and indicate that the TRPC3 channel plays a key role in the process. Taken together, the results support the view that, in macrophages, TLR4 activation increases DAG levels in the ER, increases DAG/TRPC3 colocalization, and promotes Ca^2+^ release from the organelle in a TRPC3 activitydependent manner.

**Figure 5.**
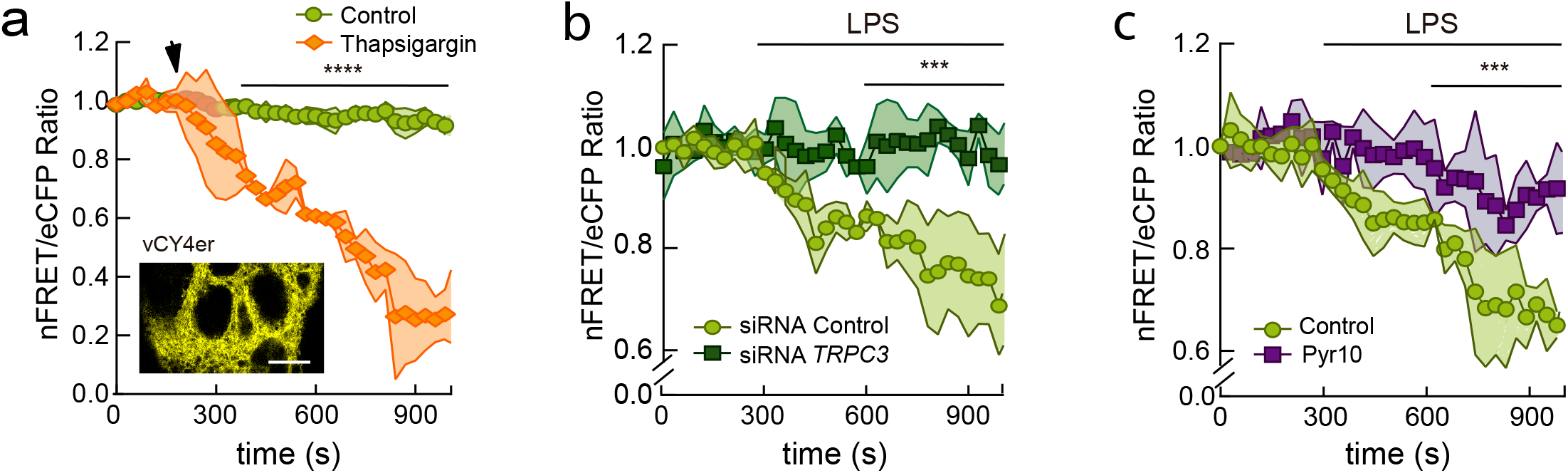
TRPC3 participates in Ca^2+^ release from the ER during LPS activation. **(a)** HEK-TLR4 cells were transfected with the vCY4er cameleon ER calcium sensor, and fluorescence from non-stimulated or thapsigargin-stimulated cells (1 μM) was analyzed by confocal microscopy. Mean nFRET/eCFP ratio ± SEM (shadow) is represented (n=40 cells). The inset shows representative vCY4er fluorescence images. Scale bar represents 10 μm. **(b)** HEK-TLR4 cells were silenced with siRNA control or against *TRPC3*, transfected with the vCY4er sensor, stimulated with 1 μg/ml LPS as indicated, and fluorescence analyzed as in e. Error (shadow) represent SEM (n=38 cells). **(c)** HEK-TLR4 cells transfected with the vCY4er sensor were stimulated with 1 μg/ml LPS as indicated, in the absence or presence of 10 μM Pyr10, and fluorescence was analyzed as in e. Error (shadow) represent SEM (n=42 cells). Multiple t-test was applied in a, b and c. ***, p < 0.001, ****, p < 0.0001.

### ER enrichment in DAG during TLR4 activation depends on lipin-1

Subsequent work was aimed to elucidate the molecular mechanisms by which DAG is generated in endomembranes during TLR4 stimulation. We have recently demonstrated that a member of the lipin family, called lipin-1, acts as a modulator of LPS-triggered signaling cascades ^20,43–45^. Hence, we considered that lipin-1 could constitute a reasonable candidate to regulate DAG levels in the ER during TLR4 challenge. As a first approach, we used propranolol, a well established inhibitor of the enzymatic activity of lipins ^46,47^, and estimated DAG production in the ER using the Daglas-em1 biosensor (**Fig. 6a**). The LPS-stimulated increase in the FRET/CFP ratio was strongly inhibited by propranolol, both in THP-1 macrophages and HEK-TLR4 cells. Moreover, use of lipin-1-deficient cells by siRNA silencing ^20^ demonstrated no significant increases in ER DAG levels during LPS challenge (**Fig. 6a)**. In addition, high colocalization between the DAG present in the ER and lipin-1 was noted, as judged by the high M1 index found between the FRET/CFP ratio from the Daglas-em1 biosensor, and the fluorescence from the chimera lipin-1-mCherry (M1= 0.72 ± 0.02) (**Fig. 6b**). A very high positive correlation (r=0.9062) was also found between the levels of DAG in the ER (FRET/CFP ratio) and lipin-1-mCherry expression levels in transfected macrophages (**Fig. 6c**). These results demonstrate that lipin-1 and DAG are in close proximity in the ER, –as would be expected from an enzyme and its product–, and that the TLR4-stimulated DAG production in the ER depends on lipin-1 expression.

**Figure 6.**
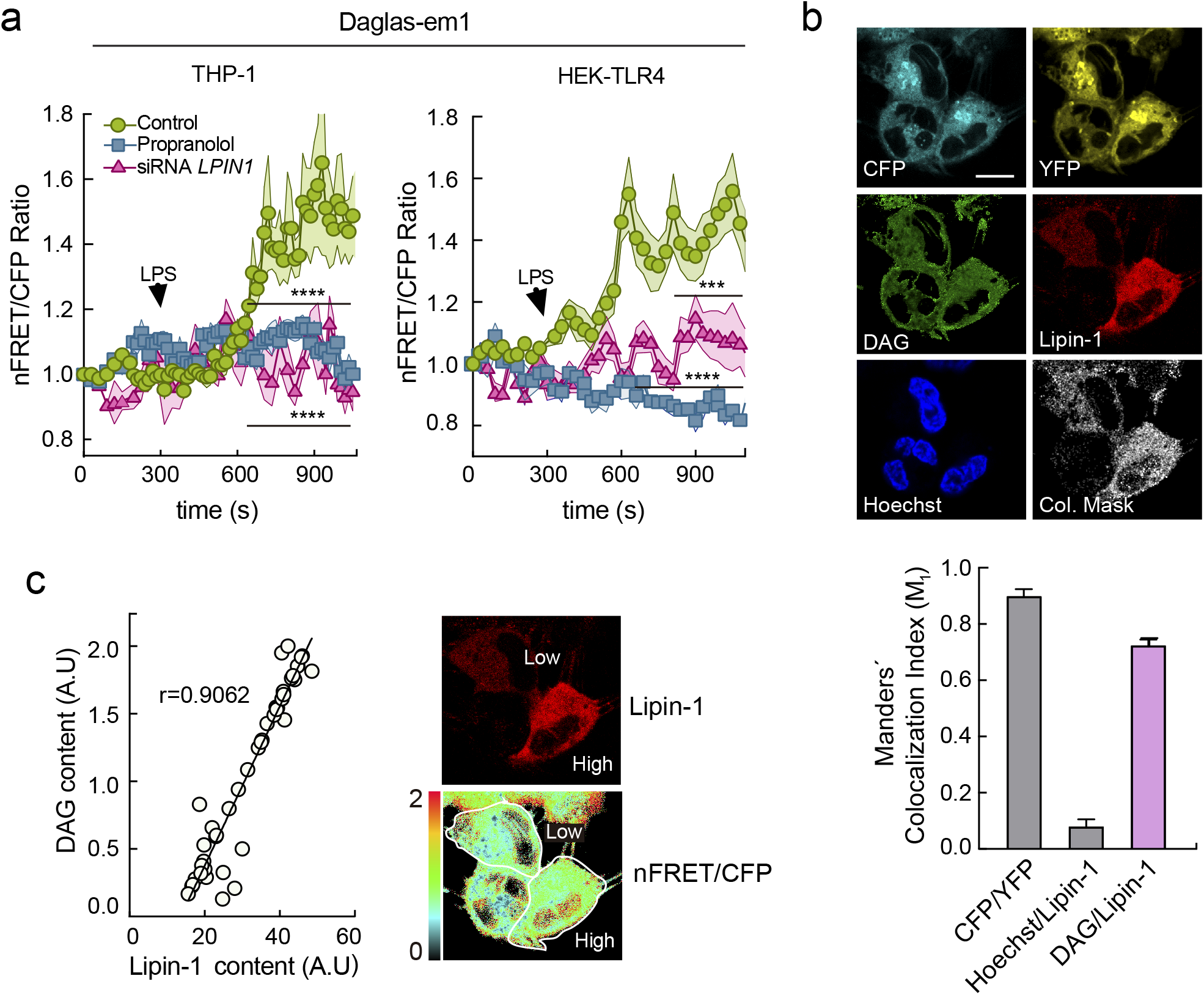
Lipin-1 mediates DAG increases in the ER during LPS activation. **(a)** THP-1 macrophages (left panel) and HEK-TLR4 cells (right panel) were transfected with the Daglas-em1 sensor. Fluorescence was analyzed by confocal microscopy before and after treating the cells with LPS (100 ng/ml and 1 μg/ml LPS respectively), in cells treated or not with 200 μM propranolol, or silenced for *LPIN1*, as indicated. Changes in nFRET/CFP ratio are shown. Multiple t-test was applied. ***, p < 0.001, ****, p < 0.0001. **(b)** THP-1 macrophages transfected with the Daglas-em1 sensor and lipin-1-mCherry were stimulated with 100 ng/ml LPS for 10 min and counterstained with Hoechst. Fluorescence from CFP (cyan), YPF (yellow), nFRET/CFP (DAG, green), lipin-1-mCherry (red) Hoechst (nuclei, blue), was analyzed by confocal microscopy and images, including merge and the colocalization mask between DAG and lipin-1-mCherry fluorescences (white), are shown (upper panel). Scale bar represents 10 μm. Manders’ colocalization indexes (m1) were calculated and are shown. Error bars represent SEM (n=22) (bottom panel). **(c)** Cells expressing lipin-1-mCherry were individually analyzed for DAG content (nFRET/CFP fluorescence) and lipin-1-mCherry content (red fluorescence) and plotted against each other to obtain a regression index (r=0.9062, n=45 cells, left panel). Representative fluorescence images are shown in the right panel.

### Lipin-1 Participates in Ca^2+^ Fluxes and TRP3-dependent Cytokine Production During TLR4 activation

If, as the previous data suggested, lipin-1 is the enzyme that provides DAG for TRPC3 activation, inhibition or absence of the enzyme should affect the TLR4-associated Ca^2+^ fluxes described above. We analyzed [Ca^2+^]i in macrophages treated with propranolol and found that the inhibitor prevented [Ca^2+^]i increases during cellular LPS treatment (**Fig. 7a**). The same effect was also observed in human macrophages silenced for *LPIN1* (**Fig. 7b**), and in murine macrophages with a spontaneous mutation in the *Lpin1* gene (*fld* animals) that eliminates lipin-1 expression ^23^ (**Fig. 7c**). Ca^2+^ responses observed in *fld* macrophages to LPS treatment were not altered by Pyr10 treatment (**Fig. 7c**), supporting the idea that lipin-1 is upstream of TRPC3 activation in this setting.

**Figure 7.**
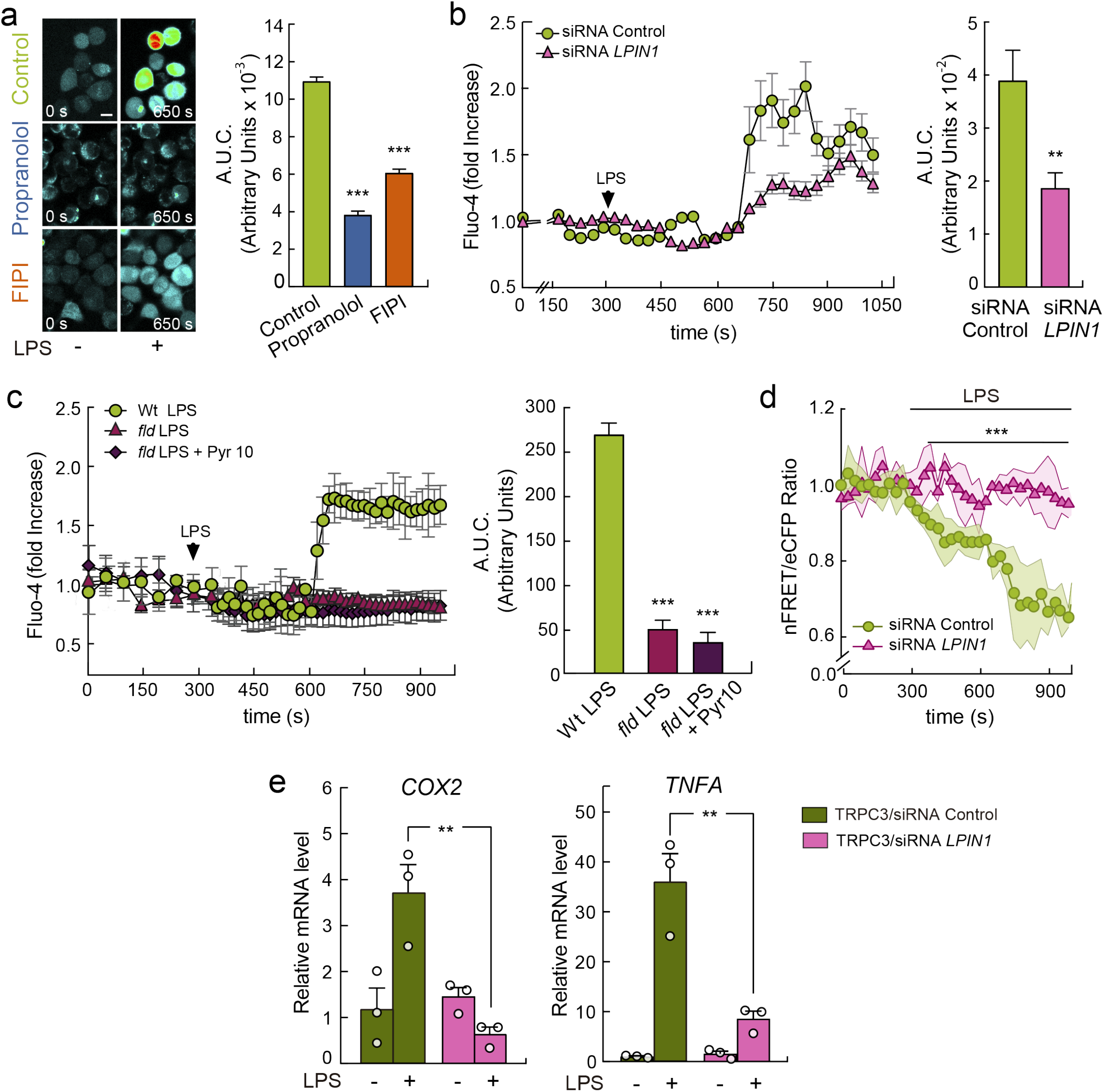
Lipin-1 participates in Ca^2+^ fluxes during LPS stimulation. **(a)** THP-1 macrophages were labeled with Fluo-4, and fluorescence was recorded before and after 100 ng/ml LPS treatment, in the absence or presence of 200 μM propranolol or 1 μM FIPI, as indicated. Representative fluorescence images (left panel) and quantification of the areas under the curves (AUC, arbitrary units; right panel) are shown. Error bars represent SEM (n= 25). ***, p < 0.001, by Student’s t test. **(b)** THP-1 macrophages silenced with siRNA control or against *LPIN1* were labeled with Fluo-4, and fluorescence was recorded before and after treating the cells with 100 ng/ml LPS, as indicated (left panel). Quantification of the area under the curve (AUC) is shown in arbitrary units (right panel). Error bars represent SEM (n=25). **, p < 0.01, Student’s t test. **(c)** Peritoneal macrophages from wt or lipin-1 deficient animals (*fld*) were used for [Ca^2+^]i quantification as in b. Error bars represent SEM (n=650). ***, p < 0.001, Student’s t test. **(d)** HEK-TLR4 cells were silenced with siRNA control, or against *LPIN1*, transfected with the vCY4er sensor, and changes in [Ca^2+^]er were analyzed before and after treating the cells with 1 μg/ml LPS, as indicated. Mean nFRET/eCFP ratio ± SEM is represented (n= 40 cells). Multiple t-test was applied. ***, p < 0.001. **(e)** HEK-TLR4 cells were transfected with wt TRPC3, silenced with siRNA control or against *LPIN1* and stimulated with 1 μg/ml LPS for 3h. mRNA levels for the indicated genes where analyzed by RT-qPCR. Error bars represent SEM (n=3). **, p < 0.01; by Student’s t test.

[Ca^2+^]er was studied next by using the calcium cameleon indicator vCY4er. The reduction in [Ca^2+^]er experienced by the cells during LPS treatment was significantly prevented in *LPIN1*-silenced cells (**Fig. 7d**). Further evidence for the implication of lipin-1 in TRPC3 activation was obtained when we assessed the upregulation of inflammatory genes in HEK-TLR4 cells that, while overexpressing TRPC3, exhibited reduced levels of lipin-1 by gene silencing. The absence of lipin-1 clearly prevented TRPC3 from being involved in LPS-induced upregulation of *COX2* and *TNFA* (**Fig. 7e**). Collectively, these results highlight a key role for lipin-1 in regulating Ca^2+^ mobilization from the ER and TRPC3-mediated inflammation during TLR4 activation. The substrate of lipins, PA, can be produced in activated cells by the direct action of phospholipase D (PLD) on membrane glycerophospholipids ^48^. We treated macrophages with the specific PLD inhibitor FIPI ^49^, and found that it affected the rise in [Ca^2+^]i during TLR4 activation (**Fig. 8a**). This finding is consistent with a scenario whereby PLD provides phosphatidic acid for DAG production and TRPC3 activation.

**Figure 8.**
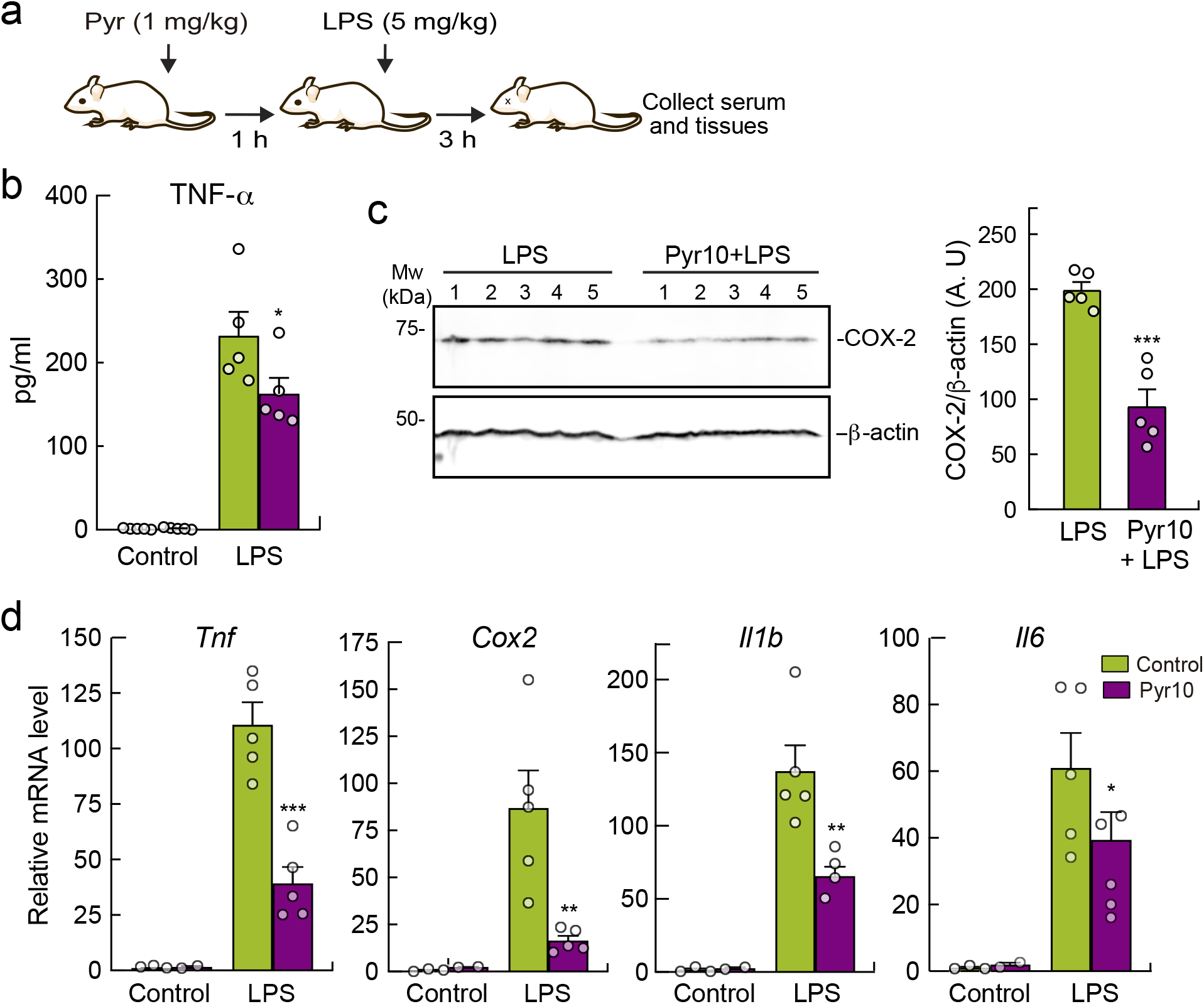
Pyr10 treatment reduces LPS-induced inflammation in mice. **(a)** Scheme of Pyr10 treatment and LPS-induced inflammation. Mice were intraperitoneally (i.p.) injected with 1 mg/Kg Pyr10. One hour later they were treated (i.p.) with 5 mg/Kg of LPS. Serum and livers were collected after three hours of treatment. **(b)** TNF-α levels present in the serum of the animals were analyzed by specific ELISA. Error bars represent SEM (n=5). **(c)** COX-2 levels were analyzed by immunoblot in liver homogenates using specific antibodies. β-Actin was used as loading control (left panel). Quantification of the bands is shown. Error bars represent SEM (n=5) (right panel). **(d)** mRNA levels of the indicated genes were analyzed in liver by RT-qPCR. Error bars represent SEM (n=5). *, p < 0.05; **, p < 0.01; ***, p<0.001, Student’s t test.

### TRPC3 Inhibition Ameliorates Systemic Inflammation *in vivo*

We evaluated the pathophysiological relevance of TRPC3 *in vivo* by analyzing the effects of the inhibitor Pyr10 during an acute systemic inflammatory response induced by treating mice with LPS (**Fig. 8**). The animals were pretreated i.p. with Pyr10 (1mg/Kg) for 1 h and then, LPS was injected i.p. (5 mg/Kg) for 3 h. Measurement of cytokine levels in blood serum showed decreased levels of TNF-α (**Fig. 8b**). COX-2 protein levels were assayed by immunoblot in liver, where LPS has a very potent effect. The results showed that the enzyme was less expressed in the Pyr10-pretreated group, implying reduced production of inflammatory eicosanoids (**Fig. 8c**) ^50^. Also, analysis of mRNA levels for different inflammatory genes showed that the drug strongly prevented the upregulation of *Cox2, Tnfa, Il1b*, and *Il6* mRNA levels in the liver (**Fig. 8d**). These results demonstrate that the inhibition of TRPC3 with Pyr10 reduces the inflammatory response induced by TLR4 challenge in animals.

## DISCUSSION

The involvement of Ca^2+^ fluxes in LPS signaling in macrophages and their relevance to mount a full inflammatory reaction have long been recognized ^6^. However, the molecular players regulating these processes are still ill defined. In the present work we describe a new key effector for LPS-triggered Ca^2+^ signaling, the ion channel TRPC3. TRPC3 participates in the release of Ca^2+^ from intracellular stores triggering a cascade of events that impact on TLR4 endocytosis, MAPK activation, NF-κB translocation to the nucleus and upregulation of inflammatory genes **(Fig. 9)**. This is relevant not only in the context of immune signaling, but also because it provides a straightforward answer to the long-standing questions about the molecular origin of the cellular DAG utilized for TRPC3 regulation ^51^, and the physiological relevance of the intracellular TRPC3 pools. Since the development of new pharmacological strategies that use TRPC3 as a target are well advanced, and we have observed that selective inhibition of TRPC3 reduces inflammation in mice, our study lays the ground for new therapies against inflammatory-based diseases.

**Figure 9.**
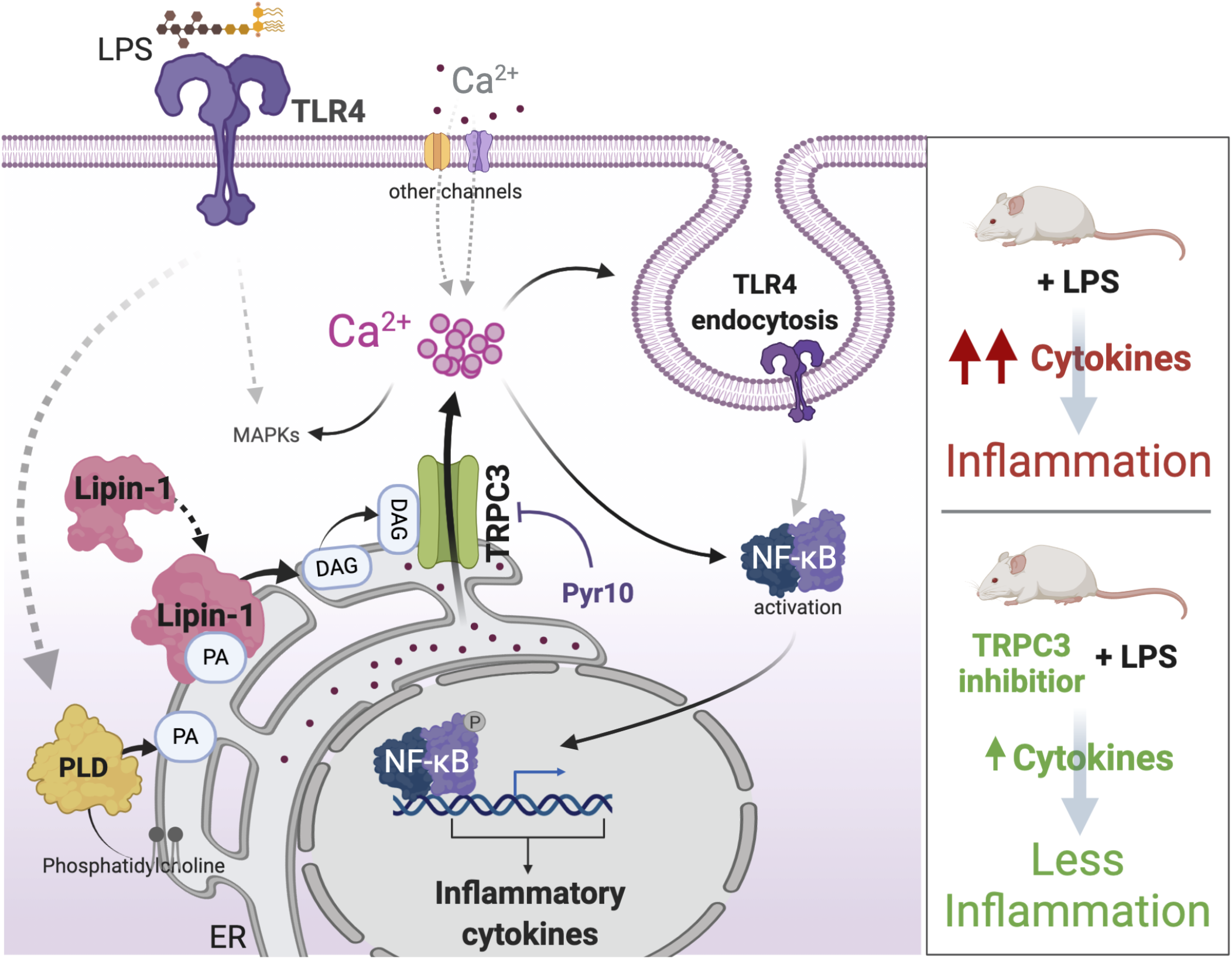
Model of TRPC3 signaling during TLR4 activation. When TLR4 senses LPS, PA generated by PLD acts as an anchor and substrate for lipin-1 at the ER. Lipin-1-derived DAG is then recognized by a pool of TRPC3 channels located in those membranes, mediating Ca^2+^ release from the organelle. The decrease in ER Ca^2+^ levels, and/or perhaps other signals that occur at the plasma membrane, stimulate then the entrance of extracellular Ca^2+^ through different plasma membrane channels. Increased levels of [Ca^2+^]i initiate a signaling cascade that ends in TLR4 endocytosis and NF-κB activation upregulating the expression of inflammatory cytokines. MAPKs seem to also be activated by Ca^2+^ fluxes dependent on TRPC3. Consequently, pharmacological inhibition of TRPC3 (Pyr10) protects mice from LPS-induced systemic inflammation.

A striking feature of TRPC3 channels is their dependence on DAG. Regulated increases in DAG levels provide an efficient means to turn on the activity of TRPC3 in accordance with the activation state of the cells ^17,18^, facilitating its role in modulating cellular Ca^2+^ fluxes. It is generally accepted that phosphoinositide-specific phospholipase C, by producing inositol 1,4,5-trisphosphate (IP_3_) and DAG, impacts as a double-edged sword on intracellular Ca^2+^ signaling ^13^. IP_3_ interacts with specific receptors in the ER favoring Ca^2+^ release, and DAG activates TRPCs at the plasma membrane, allowing Ca^2+^ entry from the extracellular medium ^13^. However, the unanticipated findings that Pyr10-sensitive ion currents are not altered by LPS at the plasma membrane level, and that TRPC3 is mainly present in intracellular membranes in macrophages, makes it difficult to envision a mechanism for channel activation coupled to phosphoinositide hydrolysis at the plasma membrane ^52^. Our results are consistent with the notion that intracellular TRPC3 is activated by DAG generated close to the channel, which allows the release of Ca^2+^ from internal stores. Considering possible candidate enzymes that could be involved in the regulation of intracellular DAG levels during TLR4 challenge, lipin-1 fulfills key requirements. For example, although lipin-1 does not reside permanently in the ER, it is able to translocate to ER membranes via a polybasic domain that facilitates binding to its PA substrate ^53^. Interestingly, lipin-1 has previously been found to regulate cellular DAG levels during macrophage activation by LPS ^19,20^ and, in this manner, impact on many cellular and biological processes of activated macrophages. However, the molecular mechanisms governing these effects were not identified. The discovery that lipin-1 regulates TRPC3 activation and associated Ca^2+^ fluxes supports the view of the phosphatase having a wider role than anticipated during macrophage activation. In fact, from a signaling point of view, lipin-1 resembles the effects triggered by phospholipase C at the plasma membrane, i.e. promoting the accumulation of DAG and initiating Ca^2+^ fluxes. Thus, our data support the view that during TLR4 activation, similar processes take place at two different cellular locations, i.e. the plasma membrane and at the ER, generating DAG to activate signaling effectors and, by different mechanisms, regulating the release of Ca^2+^ from the ER.

Consistent with the above scenario, organelle lipidomics performed in TLR4-activated macrophages has revealed that the ER undergoes a profound lipid remodeling ^54^. Interestingly, a few molecular species of PA increase in this compartment, namely PA(32:1), PA(34:1), PA(36:0), PA(40:3), and PA (40:6). It is clear that some of these species may serve as substrates for phospholipid synthesis –a process that is known to occur during TLR4 activation –, but others would participate in signaling either directly or via conversion to other signaling lipids such as DAG ^20,55^. We have shown here that inhibition of PLD activity decreases Ca^2+^ fluxes during LPS stimulation. Macrophages express PLD1 and PLD2, and both have the capacity to participate in TLR4 signaling ^56–59^. While PLD1 is present at the plasma membrane and intracellular vesicles, PLD2 seems to localize at perinuclear regions ^56^. PLD2 has an important role in sepsis models by contributing to the production of inflammatory cytokines ^60^, in a similar manner to what has been previously described for lipin-1 ^20^ or TRPC3 in this study. All of these studies support the perception that, during LPS treatment, macrophages stimulate the production of bioactive lipids, using the enzymatic machinery associated with ER membranes, which activates channels that help, or participate directly, in the release of Ca^2+^ from the organelle. These events seem to be critical for mounting an effective inflammatory response.

The participation of TRPC3 in the release of Ca^2+^ during LPS challenge does not preclude an involvement of IP_3_ receptors as well, as proposed by previous work ^7^. Interestingly, the IP_3_ antagonist utilized in those studies, (2-aminoethoxydiphenyl borate), also blocks TRPC3 ^61,62^, which makes it difficult to estimate the actual participation of IP_3_ receptors to the LPS response. Based on our studies using ER-tagged cameleon genetic probes where TRPC3 was silenced or inhibited, we estimate that TRPC3 contributes by no less than 75% of the TLR4-induced ER Ca^2+^ release. Of note, TRPC3 possesses a highly conserved IP_3_ receptor binding region in its cytoplasmic C-terminus, and in fact, TRPC3 has long been described to work in association with IP_3_ receptors in other cell systems ^16^. Future work should be directed toward determining whether TRPC3 associates with IP_3_ receptors in ER membranes and collaborates to Ca^2+^ release during LPS activation of the macrophages. Once ER Ca^2+^ levels drop in a TRPC3-dependent manner, the organelle could replace its initial Ca^2+^ concentrations thanks to other Ca^2+^ permeable channels present at the plasma membrane ^8,10,28,29,63^, contributing to the transient overall increase in [Ca^2+^]i.

To conclude, we uncover the ion channel TRPC3 as a new early player in TLR4-stimulated responses in human macrophages. Our work introduces the concept that intracellular, lipid-activated TRPC3 participates in Ca^2+^ signaling during TLR4 activation and inflammation development. Further, we have presented evidence that lipin-1 provides the DAG used for TRPC3 activation in this context. These findings help fill some gaps in our perception on how innate responses are mounted and provide new insights into the development of new therapies for inflammatory diseases.

## METHODS

### Cells

HEK293 cells expressing human TLR4/MD2/CD14 (HEK-TLR4) were generously provided by Dr. J.J. Lasarte (University of Navarra, Pamplona, Spain) were cultured in DMEM supplemented with 2 mM glutamine, 10% fetal bovine serum (FBS), 1% sodium pyruvate, 1% non-essential aminoacids (NEAA), 100 U/ml penicillin, 100 μg/ml streptomycin, 5 μg/ml blasticidin and 25 μg/ml hygromycin B at 37°C in a 5% CO_2_ humidified incubator. The cells were passaged twice a week by detachment with 1mM EDTA in PBS.

The human monocytic cell line THP-1 was maintained in RPMI 1640 medium supplemented with 10 mM HEPES, 10% FBS, 100 U/ml penicillin, 100 μg/ml streptomycin, 2 mM glutamine, 1% sodium pyruvate, 1% NEAA and 50 μM β-mercaptoethanol at 37°C in a 5% CO_2_ humidified incubator. For differentiation to a macrophage phenotype, THP-1 cells were incubated with 25 ng/ml PMA for 24 h, after which they were left to rest for 48 h.

Peritoneal macrophages were obtained, as previously described ^20^. Briefly, mouse peritoneal cavity was flushed with 5 ml ice-cold PBS. Cells were then centrifuged at 300 g for 10 min and allowed to adhere to glass coverslips overnight. Nonadherent cells were washed away with PBS and attached cells were maintained in culture until use.

### Plasmids and Mutagenesis

Cells were transfected with 1 μg pEYFP-C1-hTRPC3 (kindly provided by Dr. K. Groschner, University of Graz, Austria). Wild-type hTRPC3 was mutagenized within the transmembrane pore domain (S6 helix) in order to modify its DAG discrimination capacity ^17^ by replacing Gly^652^ with Ala (G652A) using the Quick-Change XL site-directed mutagenesis kit (Stratagene, La Jolla, CA) and the oligonucleotides described in Table 1. Mutagenesis was confirmed by sequencing. hTRPC6-YFP was a gift from Dr. C. Montell (The Johns Hopkins University School of Medicine, Baltimore, MD, USA) (Addgene plasmid #21084)^64^. The plasmids pcDNA3-Daglas-pm1 and pcDNA3-Daglas-em1 was a gift from Dr. Y. Umezawa (Department of Chemistry, School of Science, University of Tokyo, Japan). pBudCE4.1-vYC4er was kindly provided Dr. W. Graier (University of Graz, Austria). Construct expressing lipin-1-mCherry was amplified by PCR adding 5’Xba-I and Sal-I 3’specific restriction sites and inserted into lentiviral vector pCDH-CMV-MCS-EF1α-copGFP (System Biosciences) after removing an internal Sal-I restriction site by mutagenesis. Daglas-em1 was amplified by PCR adding 5’Xba-I and Sal-I 3’specific restriction sites and inserted into lentiviral vector pCDH-CMV-MCS-EF1α-copGFP. Plasmid delivery in HEK-TLR4 cells was done using linear polyethyleneimine (PEI) 25K (Polysciences) complexed with DNA in a 2:1 ratio respectively for 24 h.

### Gene Silencing

To knock-down *TRPC3* and *LPIN1* in HEK-TLR4 cells, specific siRNAs (Sigma) were introduced into the cells using Lipofectamine RNAiMAX (Thermo Fisher Scientific) as specified by the manufacturer, and experiments were conducted 48 h later (see Table 1). For gene silencing in THP-1 macrophages, PMA was used to differentiate human THP-1 monocytes for 48 h into macrophages prior to transfection with siRNA. For transfection, macrophages were detached enzymatically by a 30 min Accutase (Gibco) I treatment. Transfection of siRNAs was then performed using a Nucleofector device ^65^ Differentiation was allowed to continue for another 48 h.

### Lentiviral Transduction

For lentiviral transduction, VSV-G–pseudotyped lentiviral particles were produced by cotransfection of 293FT cells with transfer constructs and the compatible packaging plasmids pMD2.G and psPAX2. Transfections were carried out using PEI 25K, and viruses were harvested at 48 and 72 hours after transfection. Concentrated lentiviral supernatants were used to transduce cells in the presence of 8 μg/ml Polybrene (Sigma-Aldrich), and infected cells were selected by FACS sorting and subjected to imaging experiments.

### Cytosolic Ca^2+^ Measurement

Cells were loaded with 3 μM Fluo-4-AM for 20 min in culture medium at 37°C in a 5% CO_2_ incubator. Cells were then washed in indicator-free medium to remove any dye that was nonspecifically associated with the cell surface, and then incubated for a further 20-min period to allow complete de-esterification of intracellular AM esters. Live cell fluorescence was monitored by confocal microscopy (TCS SP5X, Leica) using 488 nm laser excitation and an emission window of 500-560 nm, with the iris fully open. Before imaging started, medium was replaced by HBSS supplemented with 10 mM HEPES, with 1.3 mM CaCl_2_ and 1.3 mM MgCl_2_. We recorded a 5 min baseline before adding LPS acquiring images each 15 s. In some experiments calibration of Ca^2+^ levels was achieved at the end of each experiment as previously described by Kao *et al*. ^66^. Cells were treated with MnCl_2_ at a final concentration of 2 mM in the presence of 2.5 μM A23187, and then cells were lysed with 0.05% digitonin to obtain the fluorescence background signal. Fluorescence data were analyzed using a combination routine of ImageJ and Cell Profiler semiautomated custom pipeline.

### Live cell FRET imaging

Cells expressing FRET-based sensors (Daglas or vCY4er) were plated together with a small proportion of single-color control expressing cells on glass-bottom culture dishes (MatTek) coated with poly-L-lysine (Sigma) when improved adhesion was required. The culture medium was replaced with HBSS supplemented with 10 mM HEPES, pH 7.4, 1.3 mM CaCl_2_ and 1.3 mM MgCl_2_ for confocal imaging. Cells were imaged at 37°C on a Leica confocal system TCS SP5X, correct membrane/endomembranes localization of the Daglas sensor was confirmed by YFP imaging. For ratio imaging, three images were acquired at each time point: the donor image (*I_CFP_*) was obtained by exciting CFP (458 nm) and monitoring its emission (465-510 nm), the acceptor image (*I_YFP_*) was acquired by exciting YFP fluorescent protein (514 nm) and monitoring its emission (525-580 nm), and the FRET image (*I_FRET_*) was obtained by exciting the donor CFP (454 nm), and monitoring the emission of the acceptor YFP (525-580 nm). *I_CFP_* and *I_FRET_* where simultaneously acquired while *I_YFP_* was acquired alternately. ImageJ software with customized plugins was used to correct the background from raw images and to create ratio images and export raw data to Microsoft Excel for further ratio and correction calculations. The FRET/CFP ratio values were divided by single color YFP expressing cells values in order to correct for any laser power illumination inconsistency ^67^, then divided by the baseline normalized FRET/CFP ratio according to equation:

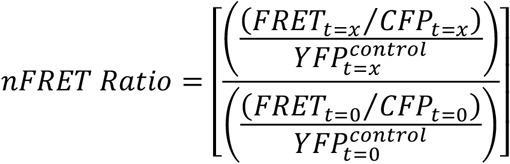

### Patch Clamp Electrophysiology

Mouse peritoneal macrophages were plated on 12-mm coverslips, previously treated with poly-L-lysine (0.01 mg/ml), for 30 min. Patch micropipettes were made from borosilicate glass (2.0 mm OD; WPI) and double pulled (Narishige) to resistances ranging from 5 to 10 MΩ when filled with the internal solution containing (in mM): 125 KCl, 4 MgCl_2_, 10 HEPES, 10 EGTA, 5 MgATP, pH 7.2 with KOH. Cells were bathed in an external solution containing (in mM): 141 NaCl, 4.7 KCl, 1.2 MgCl_2_, 1.8 CaCl_2_, 10 glucose, and 10 HEPES, pH 7.4 with NaOH. Immediately after starting the recording of whole-cell currents, the external solution was switched to a solution containing (in mM): 131 NaCl, 10 TEACl, 4.7 KCl, 1.2 MgCl_2_, 1.8 CaCl_2_, 10 glucose, 100 μM BaCl_2_, 100 μM Niflumic acid and 10 HEPES, pH 7.4 with NaOH. This solution allows to record in isolation currents mediated by TRP like channels, blocking currents through voltage-dependent K^+^ channels, inward rectifier channels and chloride channels. Cells were voltage-clamped at −40 mV. Ionic currents were recorded at room temperature using the whole-cell configuration of the patch-clamp technique. Current–voltage relationships were obtained with a 1s-ramp protocol from −100 mV to +100 mV applied every 10 s. Whole-cell currents were recorded using an Axopatch 200A patch-clamp amplifier (Axon Instruments) filtered at 2 kHz (−3 dB, 4-pole Bessel filter) and sampled at 10 kHz. Recordings were digitized with a Digidata 1200, driven by CLAMPEX 10.2 software (Molecular Devices). Electrophysiological data analyses were performed using Clampfit subroutine of the pClamp software (Axon Instruments) and with Origin 7 software (OriginLab Corporation). Current amplitude was corrected for cell size variations and expressed as current density (pA/pF), by dividing for cell capacitance values.

### Semi Quantitative PCR and RT-qPCR

Total RNA was extracted using TRIzol reagent (Ambion). cDNA was obtained using Verso cDNA kit Reverse Transcription for RT-PCR (Thermo Fisher Scientific), following the manufacturer’s instructions. For semiquantitative PCR forward and reverse primers specific to human TRPC3, 6, 7 and β-actin were used (see table). PCR reactions were carried out on an Eppendorf Mastercycler Personal thermal cycler using Taq polymerase (BioTools) and the following parameters: denaturation at 94°C for 30 s, annealing at 58°C for 30 s, and extension at 72°C for 40 s. A total of 40 cycles were performed followed by a final extension at 72°C for 5 min. PCR products were analyzed by electrophoresis with 1.5% agarose gel and visualized by GelRed staining. Bands were intensity quantified with Image Studio Software (LiCor). Quantitative real time-PCR analysis was performed in an ABI 7500 (Applied Biosystems) using specific primers, 20 ng cDNA and the Brilliant III Ultra-Fast SYBR Green qPCR Master Mix (Agilent Technologies). Relative mRNA expression was obtained using the ^ΔΔ^Ct method using β-actin as reference gene ^68^.

### Colocalization analysis

Confocal z-stacks images were deconvolved using an ImageJ parallel iterative plugin (Wiener Filter Preconditioned Landweber (WPL) method) after calculating the experimental setup point spread function (PSF) using sub resolution (0.17 μm) fluorescent beads. Colocalization was performed with JACoP plugin in ImageJ. Background levels were obtained by measuring the mean intensity of each signal outside the cells and were subtracted; negative pixel values were clipped to zero. Positive values were selected by Costes automatic thresholding, removing the bias of visual interpretation ^69^. Colocalization index Manders’ *m1* and *m2*, were calculated ^40^.

Manders’ split coefficients are based on the Pearson’s correlation coefficient but avoid issues relating to absolute intensities of the signal, since they are normalized to total pixel intensity ^40^. These coefficients vary from 0 (non-overlaping) to 1 (100% colocalization). The index m1 is defined as the percentage of above-background pixels from the first channel (green) that overlap above-background pixels from the second channel (red). This index is sensitive to changes in the background but not to differences in the intensity of overlapped pixels and is suitable to apply in images with a high and clear signal to background ratio.

### Immunocytochemistry analysis

For intracellular staining of endogenous TRPC3 or NFkB-p65, cells were fixed with 4% paraformaldehyde for 20 min in LabTek II chambers and then permeabilized with 0.1% Triton X-100 for 20 min at RT. Cells were treated with 5% goat serum for 30 min and incubated with antibodies against TRPC3 (Alomone Labs) or NFkB-p65 (Cell Signaling) for 1h. After washing, cells were incubated with spectrally appropriate Alexa Fluor-conjugated Fab fragments against primary species antibodies (Molecular Probes) for 1h. After intracellular staining, nuclei were counterstained with the DNA binding dye Hoechst 33342 (Invitrogen). All images were captured with a Leica confocal system TCS SP5X inverted microscope. Leica Application Suite Advanced Fluorescence software was used for the capture, and ImageJ was used for deconvolution and image presentation. Translocation was analyzed using Cell Profiler software (Broad Institute) with a custom pipeline (available upon request) that automatically detects cell boundaries and calculates the ratio between cytoplasm and nuclei fluorescence median.

### TLR4 endocytosis by Flow Cytometry

For these experiments, the procedure described by Schappe et al.^70^ was followed, with minor modification. Briefly, after LPS stimulation, the cells were washed with cold PBS and collected at 4 °C in FACS buffer (PBS supplemented with 0.5% BSA and 2 mM EDTA). After Fc receptor blocking, cells were stained with a monoclonal antibody against TLR4 (clone HTA125) conjugated with Phycoerythrin PE for 30 min following the manufacturer instructions (Invitrogen). Stained cells were washed in cold FACS buffer, resuspended in 500 μl FACS buffer, and kept on ice for immediate analysis. Mean fluorescence intensity (MFI) of TLR4 was measured from unstimulated and LPS stimulated cells using a Beckman Coulter Gallios flow cytometer. Relative percentage of surface TLR4 at given timepoints was represented as:

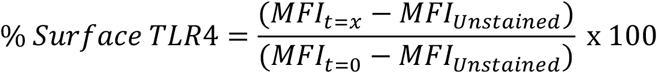

### Animals

BALB/cJ mice were obtained from the University of Valladolid Animal House (10-12-week-old males). The mice were intraperitoneally injected with the TRPC3 specific inhibitor Pyr10 (1 mg/Kg) for 1h and then treated with LPS at a sublethal dose of 5 mg/Kg (i.p.). For the analysis of proinflammatory factors, animals were sacrificed by ketamine (100 mg/Kg): xylacine (10 mg/Kg) administration and cervical dislocation 3 h after LPS treatment. Blood was collected through cardiac puncture. Livers were collected in RNAlater (Ambion) for further analysis by real-time PCR or in protein lysis buffer for western blot analysis. Serum from animals were used for quantification of ITNF-α by specific ELISAs (Thermo) following the manufacturer’s instructions. BALB/cByJ-Lpin1fld/J mice carrying a spontaneous mutation in the *Lpin1gene* (fatty liver dystrophy, *fld*) were also used ^20^. All the protocols and procedures were approved by the Institutional Animal Care and Usage Committee of the University of Valladolid and are in accordance with the Spanish and European Union guidelines for the use of experimental animals.

## Quantification and Statistical Analysis

Statistical details of experiments are indicated in the figure legends. All data analyses were performed with Prism software (GraphPad). All data are presented as means ± standard error of the mean (SEM), indicating individual biological replicates. No statistical analysis was used to predetermine sample size. For *in vivo* experiments, animals were randomized in different treatment groups. All datasets were analyzed by unpaired two-tailed Student’s t test. Welch’s correction was performed when the variances were significantly different. Some data required multiple t-test, with a false discovery rate of 1 % based on two stage step up by Benjamini, Krieger and Yekutieli.

## ACKNOWLEDGMENTS

We thank Montserrat Duque for technical assistance and Lorena García for her critical reading. This work was supported by the Spanish Ministry of Economy, Industry, and Competitiveness (grant SAF2016-80883-R) and the Spanish Ministry of Science and Innovation (PID2019-105989RB-I00). Centro de Investigación Biomédica en Red de Diabetes y Enfermedades Metabólicas (CIBERDEM) is an initiative of Instituto de Salud Carlos III. The authors declare no competing financial interests.

## AUTHOR CONTRIBUTIONS

Conceptualization, J.C., and M.A.B.; Methodology, J.C., C.M., J.R.L-L., and M.A.B.; Investigation, J.C., C.M., and J.R.L-L.; Writing – Original Draft, J.C.; Visualization, J.C., M.A.B.; Supervision, J.B., and M.A.B.; Writing – Review & Editing, J.C., J.B., and M.A.B; Funding Acquisition, J.B., and M.A.B.

## MATERIALS & CORRESPONDENCE

* Correspondence should be addressed to J.C or M.A.B. Further information and request for resources and reagents should be directed to and will be fulfilled by the lead contact, Prof. María A. Balboa.

