## Supplemental Figure 1 for "Intracellular pools of DAG-activated TRPC3 channels are essential for TLR4 activation"

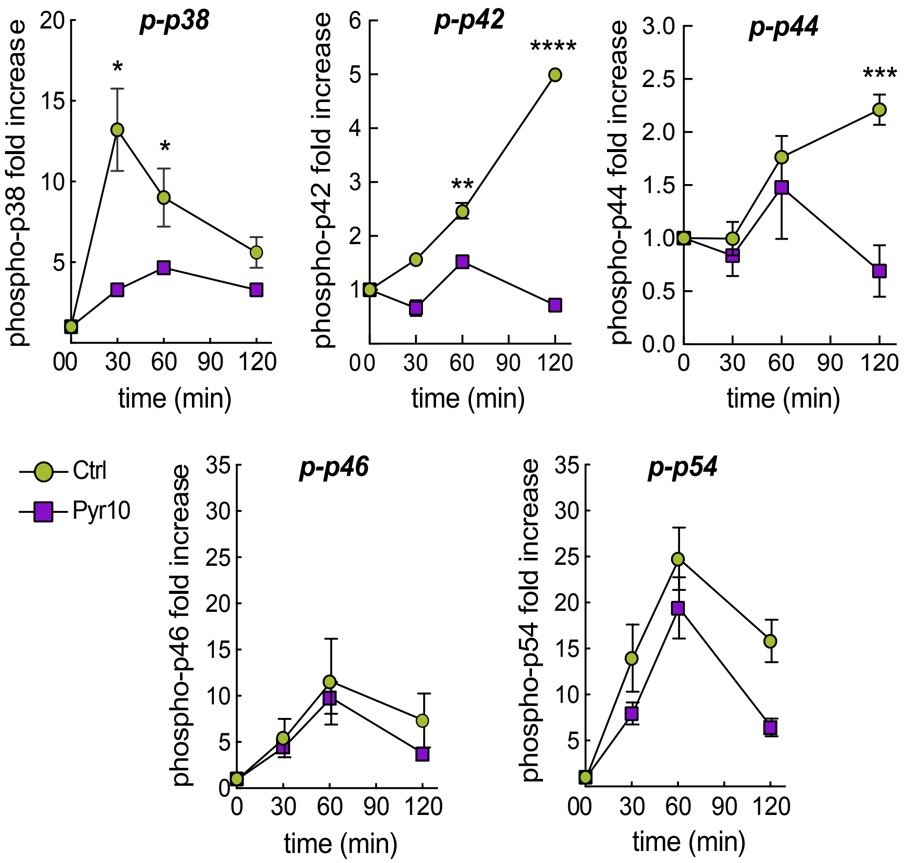


**Figure S1 (related to Fig. 1). TRPC3 is important for MAPKs activation during LPS activation.** THP-1 macrophages were pretreated with 10 μM Pyr10 for 30 min and then treated with 100 ng/ml LPS for the indicated periods of time. Proteins were evaluated by immunoblot using anti-phospho-specific antibodies against the MAPKs p38, ERK (p42/p44) and JNK (p46/p54). β-actin was used as the loading control. Quantification of phosphoproteins relative to time 0 is represented. Error bars represent SEM (n=3). *, $p < 0.05$; **, $p < 0.01$; ***, $p < 0.001$, by Student's t test.
