## Supplemental Information for "Intracellular pools of DAG-activated TRPC3 channels are essential for TLR4 activation"

### Lipin-1-derived Diacylglycerol Activates Intracellular TRPC3 Which is Critical for TLR4-induced $Ca^{2+}$ Fluxes and Inflammation

Javier Casas, Clara Meana, José Ramón López-López, Jesús Balsinde, and María A. Balboa.

#### *MATERIALS AND METHODS*

**Immunoblots.**

Cells or tissues were lysed in a buffer 1% Triton X-100, 50 mM Tris, pH 7.4, 150 mM NaCl, 1 mM EDTA, 1 mM EGTA, 5 mM $Na_4P_2O_7$, 50 mM sodium β-glicerol phosphate, 270 mM sucrose, 0.1% β-mecaptoethanol, 1 mM $Na_3VO_4$, 10 mM NaF, 1 mM phenylmethyl sulphonyl fluoride and a protease inhibitor 'cocktail' (Sigma). Cell proteins were separated by 10% reducing SDS–polyacrylamide gel electrophoresis and transferred to nitrocellulose membrane. The membranes were incubated with specific primary antibodies followed by incubation with goat anti-rabbit IgG IR-Dye 680RD-conjugated (LiCor) or goat anti-mouse IgG IR-Dye 800CW-conjugated (LiCor) and detected and accurately quantified with a LiCor Odyssey Fc infrared imaging system. For sequential primary antibodies incubations, membranes were stripped with NaOH 0.2 % for 20 min before re-blocking.

#### *FIGURE*

**Figure S1 (related to Fig. 1).** TRPC3 is important for MAPKs activation during LPS activation. THP-1 macrophages were pretreated with 10 μM Pyr10 for 30 min and then treated with 100 ng/ml LPS for the indicated periods of time. Proteins were evaluated by immunoblot using anti-phospho-specific antibodies against the MAPKs p38, ERK (p42/p44) and JNK (p46/p54). β-actin was used as the loading control. Quantification of phosphoproteins relative to time 0 is represented. Error bars represent SEM (n=3). *, $p < 0.05$; **, $p < 0.01$; ***, $p < 0.001$, by Student's t test.

***Supplementary Resources Table***

| Mutagenesis primers | Sequence |
|---|---|
| G652A_fwd | 5′-GAAAATATTGGATACGTTCTTTATGCAATATACAATGTAACTATGGTGGTC-3′ |
| G652A_rev | 5′-GACCACCATAGTTACATTGTATATTGCATAAAGAACGTATCCAATATTTTC-3′ |
| **siRNA** | **Sequence** |
| *TRPC3*_s | 5′-GCUCUUGACGAUCUGGUAUGA[dT][dT]-3′ |
| *TRPC3*_as | 5′-UCAUACCAGAUCGUCAAGAGC[dT][dT]-3′ |
| *LPIN1*_s | 5′- GGAGUGUCUUUGAAUAGAA[dT][dT]-3′ |
| *LPIN1*_as | 5′- UUCUAUUCAAAGACACUCC[dT][dA] - 3′ |
| **Standard PCR primers** | **Sequence** |
| *TRPC3*_fwd | 5′- GGAAAAACATTACCTCCACCTTTCA -3′ |
| *TRPC3*_rev | 5′- CTCAGTTGCTTGGCTCTTGTCTTCC -3′ |
| *TRPC6*_fwd | 5′-AAGACATCTTCAAGTTCATGGTC-3′ |
| *TRPC6*_rev | 5′-TCAGCGTCATCCTCAATTTCC-3′ |
| *TRPC7*_fwd | 5′- TGGGTTGTATTTGGCACCTC-3′ |
| *TRPC7*_rev | 5′- TGGGTTGTATTTGGCACCTC-3′ |
| *ACTB*_fwd | 5′- CAGAGCAAGAGAGGCATCCT-3′ |
| *ACTB*_rev | 5′- ACGTACATGGCTGGGGTG-3′ |
| **qRT-PCR primers** | **Sequence** |
| *TRPC3*_fwd | 5′- AGAATGACTATCGGAAGCTCTCC -3′ |
| *TRPC3*_rev | 5′- GGCAAGTTTGACACGACTTAATG -3′ |
| *TRPC6*_fwd | 5′- GTGATCGCTCCACAAGCCTAT -3′ |
| *TRPC6*_rev | 5′- CTGCCAACTGTAGGGCATTCT -3′ |
| *ACTB*_fwd | 5'-ATTGCCGACAGGATGCAGAA-3' |
| *ACTB*_rev | 5'-GCTGATCCACATCTGCTGGAA-3' |
| *COX2*_fwd | 5'-GTGCAACACTTGAGTGGCTAT-3' |
| *COX2*_rev | 5'-AGCAATTTGCCTGGTGAATGAT-3' |

| | |
|---|---|
| *TNFA*_fwd | 5'-ATGAGCACTGAAAGCATGATCC-3' |
| *TNFA*_rev | 5'-GAGGGCTGATTAGAGAGAGGTC-3' |
| *IL1B*_fwd | 5'-ATGATGGCTTATTACAGTGGCAA-3' |
| *IL1B*_rev | 5'-GTCGGAGATTCGTAGCTGGA-3' |
| *IL6*_fwd | 5'-AAATTCGGTACATCCTCGACGG-3' |
| *IL6*_rev | 5'-GGAAGGTTCAGGTTGTTTTCT -3' |
| *IL12B*_fwd | 5'-CAGCAGTTGGTCATCTCTTGG-3' |
| *IL12B*_rev | 5'-GGTCCAGGTGATACCATCTTCT-3' |
| *IL23A*_fwd | 5'-GCCTTCTCTGCTCCCTGATA-3' |
| *IL23A*_rev | 5'-GACTGAGGCTTGGAATCTGC-3' |
| *Cox2*_fwd | 5'-TGAGCAACTATTCCAAACCAGC-3' |
| *Cox2*_rev | 5'-GCACGTAGTCTTCGATCACTATC-3' |
| *Tnfa*_fwd | 5'-ACGGCATGGATCTCAAAGAC-3' |
| *Tnfa*_rev | 5'-AGATAGCAAATCGGCTGACG-3' |
| *Il1b*_fwd | 5'-GCAACTGTTCCTGAACTCAACT-3' |
| *Il1b*_rev | 5'-ATCTTTTGGGGTCCGTCAACT-3' |
| *Il6*_fwd | 5'-TAGTCCTTCCTACCCCAATTTCC-3' |
| *Il6*_rev | 5'-TTGGTCCTTAGCCACTCCTTC-3' |
| *Gapdh*_fwd | 5'-AGGTCGGTGTGAACGGATTTG-3' |
| *Gapdh*_rev | 5'-TGTAGACCATGTAGTTGAGGTCA-3' |

| **Antibodies** | **Source** | **Identifier/RRID** |
|---|---|---|
| Rabbit Monoclonal NF-κB p65 (clone D14E12) XP | Cell Signaling Technologies | #8242/AB_10859369 |
| Rabbit polyclonal TRPC3 | Alomone Labs | #ACC-016/AB_2040236 |
| F(ab')2-Goat anti-Rabbit IgG (H+L) 2[a] Antibody, Alexa Fluor 488 | Thermo Scientific | A-11070/AB_142134 |
| F(ab')2-Goat anti-Rabbit IgG (H+L) 2[a] Antibody, Alexa Fluor 594 | Thermo Scientific | A-11072/AB_142057 |
| PE anti-human CD284 (TLR4) (clone HTA125) | BioLegend | #312805/AB_314954 |

| | | |
|---|---|---|
| Rabbit Monoclonal Phospho-p38 MAPK (D3F9) XP | Cell Signaling Technologies | #4511/AB_2139682 |
| Rabbit Polyclonal Phospho-p44/42 MAPK (Erk1/2) | Cell Signaling Technologies | #9101/AB_331646 |
| Rabbit Monoclonal Phospho-SAPK/JNK (81E11) | Cell Signaling Technologies | #4668/AB_823588 |
| Mouse Monoclonal Anti-β-Actin (Clone: AC15) | Sigma-Aldrich | #A5441/AB_476744 |
| Goat anti-mouse IgG IRDye 800CW | Li-Cor | #925-32210/AB_2687825 |
| Goat anti-rabbit IgG IRDye 680RD | Li-Cor | #925-68071/AB_2721181 |
| TNF alpha Mouse Uncoated ELISA Kit | Thermo Scientific | #88-7324-22/AB_2575076 |
| **Reagents** | **Source** | **Identifier** |
| *E. coli* Lipopolysaccharides O111:B4 | Sigma-Aldrich | L2630 |
| PYR10 | Sigma-Aldrich | SML1243 |
| 1-Oleoyl-2-acetyl-*sn*-glycerol (OAG) | Sigma-Aldrich | #O6754 |
| FIPI | Cayman Chemical | #13563 |
| Propranolol | Sigma-Aldrich | #P0884 |
| A23187 | Sigma-Aldrich | #52665-69-7 |
| Thapsigargin | Sigma-Aldrich | #67526-95-8 |
| PMA | Sigma-Aldrich | #16561-29-8 |
| Hoechst 33342 | Invitrogen | #H3570 |
| Fluo-4 AM | Invitrogen | #F14201 |

***Table 1***
